# A Deep Learning approach for time-consistent cell cycle phase prediction from microscopy data

**DOI:** 10.1101/2025.05.16.654306

**Authors:** Thomas Bonte, Oriane Pourcelot, Adham Safieddine, Floric Slimani, Florian Mueller, Dominique Weil, Edouard Bertrand, Thomas Walter

## Abstract

The cell cycle consists of four phases and impacts most cellular processes. In imaging assays, the cycle phase can be identified using dedicated cell-cycle markers. However, such markers occupy fluorescent channels that may be needed for other reporters. Here, we propose to address this limitation by inferring the phase from a widely used fluorescent reporter: SiR-DNA. Our method is based on a variational auto-encoder, enhanced with two auxiliary tasks: predicting the intensity of phase-specific markers and enforcing the latent space temporal consistency. Our model is freely available, along with a new dataset comprising over 600,000 annotated HeLa Kyoto nuclear images.

## Background

The cell cycle is a tightly regulated series of stages resulting in cell growth, DNA replication, organelle duplication, and the partitioning of cellular components through cell division. The cell cycle consists of four distinct phases: G1 (gap 1), S (synthesis), G2 (gap 2), and M (mitosis), each characterized by specific gene expression programs. Throughout these phases, cells dynamically regulate gene expression, protein synthesis, and intracellular organization to ensure proper division and function. Many aspects of a cell depend directly or indirectly on its cell cycle phase, including gene expression [1], transcriptional activity and histone modification [2], protein interaction states [3] and drug sensitivity [4, 5]. Therefore, interpreting observations, such as localization patterns of RNAs and proteins or cellular phenotypes in the context of the cell cycle phase has the potential to reveal important functional dependencies that might otherwise remain hidden.

Cellular imaging has been widely used to investigate the cell cycle, often relying on fluorescent markers such as FUCCI, whose expression levels are correlated with the cell cycle phase [6–9]. However, such markers are invasive and may introduce experimental biases [10, 11]. Moreover, they occupy fluorescent channels that might be essential for other readouts when the primary focus is not the cell cycle itself. For instance, in studies of RNA [12] or protein localization [13], it may be important to examine how localization patterns depend on the cell cycle. In these cases, reserving two fluorescent channels for cell cycle markers reduces the ability to simultaneously capture other relevant signals. This limitation is particularly problematic when cell cycle information is needed as a complementary feature in assays focused on other biological processes.

Deep learning, and representation learning in particular, has demonstrated remarkable efficiency in extracting subtle cellular morphological features to significantly advance our understanding of biological processes [14–21]. Representation learning is a self-supervised learning approach designed to capture rich representations directly from data without prior assumptions or need for annotations. It extracts meaningful features from cellular phenotypes, which can then be used for various downstream tasks both at single-cell and subcellular resolutions: clustering [16, 18, 21], cell state classification [15, 17], cell cycle phase classification [19, 20], subcellular protein localization [18, 19].

Several recent studies have proposed deep and/or representation learning strategies for performing cell cycle phase classification without dedicated cycle markers. Deep Convolutional Neural Networks (CNN) have been proposed for cell cycle phase classification, learning complex patterns from labeled microscopy images in a fully supervised manner [22–25], operating either on label-free or fluorescence microscopy. Alternatively, meaningful hand-crafted features can be extracted from microscopy data for classification using Support Vector Machine (SVM) [25, 26]. Another approach is self-supervised learning which reduces the need for large training datasets while enabling more generalizable and robust feature extraction [19, 20]. The extracted features can then be used to train a (small) classification network to predict the cell cycle phases. Finally, [27] uses a U-Net model to directly predict a cell cycle mask.

However, the majority of these approaches simplify the classification task by merging the three distinct cell cycle phases G1, S, and G2/M. These phases are either grouped together into a single class, referred to as *interphase* [19, 22] or split into two broader categories for binary classification: G1/S vs G2/M [24] or G1 vs S/G2/M [25, 26]. Such an approach overlooks the finer distinctions between the phases, which are crucial for accurately understanding cell cycle dynamics. Among the works that do not merge G1, S and G2/M, [23] proposes a method to reconstruct a cyclic latent space based on the classification of four manually defined virtual phases. However, this approach struggles to accurately distinguish between the G1, S, and G2/M phases, limiting its effectiveness for precise cell cycle phase classification. [27] presents a method for cell cycle phase classification using Spatial Light Interference Microscopy (SLIM, [28]) images. Although the approach achieves decent classification accuracies, SLIM is not widely used, which limits the general applicability of the method. Finally, [20] employs a VQ-VAE combined with Dynamic Time Warping (DTW) to classify G1, S, G2, and M phases separately. While the approach achieves a classification accuracy similar to [27], it requires live-cell imaging data, as DTW is used in conjunction with deep learning. Consequently, it does not provide a solution for standard fixed-cell microscopy data.

Therefore, there is a need for a robust cell cycle phase classification method that relies solely on widely used microscopy techniques and does not require tracking cell trajectories. In this work, we propose to leverage representation learning to determine the cell cycle phase from a DNA marker. Fluorescent DNA markers include DAPI, Hoechst or SiR-DNA, and are very commonly used fluorescent markers that may contain key information to infer the cell cycle phase of a given cell. Indeed, during the S phase, DNA replication doubles the chromatin content, leading to an increase in nuclear size and signal intensities of DNA markers [29, 30]. These changes, along with chromatin structure patterns, may serve as cell cycle-dependent features that traditional analyses might have had difficulties to leverage.

Here, we introduce Cell Cycle Variational Auto-Encoder (CC-VAE), a new self-supervised strategy based on VAE [31] to learn a meaningful latent representation of nuclei SiR-DNA [32] images and accurately perform cell cycle phase classification.

## Results

### Dataset

We introduce a new image dataset, which comprises 982,332 images of HeLa Kyoto nuclei including 636,304 labeled into one of the three classes G1, S or G2/M. Images were acquired with a 63× objective microscope. Each image contains five focal planes and three channels: a DNA marker (SiR-DNA) and two PIP-FUCCI markers derived from the PIP-FUCCI system [9]: mCherry-Gem_1-110_ and PIP-mVenus. These markers allow for the inference of the cell cycle phase for each individual cell.

We performed time-lapse microscopy via live-cell fluorescence imaging of HeLa Kyoto cells labeled with SiR-DNA and the two PIP-FUCCI markers. Images were captured every 20 minutes over a period of 40 hours. While this temporal resolution is crucial for confidently assessing the cell cycle phase for each cell, it is important to note that our classification method is designed to process a single image as input, here the DNA-labeled cells. Cell cycle annotation was based on the temporal evolution of the PIP-FUCCI intensities across their area. By doing this, the cell cycle phase was determined for 636,304 nuclei. 346,028 nuclei remained unlabeled due to the inability to reliably determine cell cycle phase boundaries based on their PIP-FUCCI intensity changes, as the tracks were either too short, displayed only one of the three phases, or were truncated at the end of the video.

Fig.1 displays some example nuclei from our annotated dataset. G1 nuclei express only PIP-mVenus fluorescence, S nuclei express only mCherry-Gem_1-110_ fluorescence, and G2 nuclei express both signals [9].

**Fig. 1:**
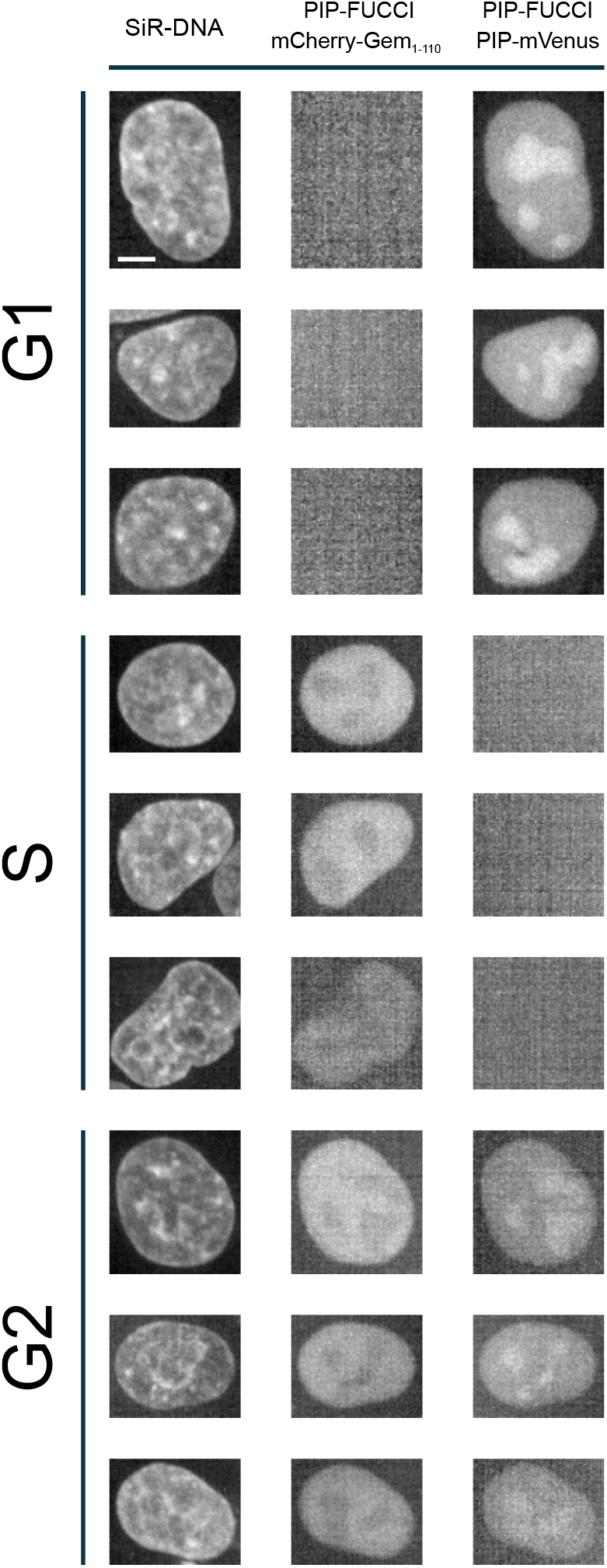
Example nuclei from our annotated dataset. G1 nuclei express only PIP-mVenus fluorescence, S nuclei express only mCherry-Gem_1-110_ fluorescence, and G2 nuclei express both signals. Scale bar: 5µm.

### Overview of CC-VAE

Our method, CC-VAE, is based on a customized *β*-Variational Auto-Encoder (*β*-VAE, [33]) architecture to learn an unsupervised latent representation of nucleus images (see Fig.2a).

**Fig. 2:**
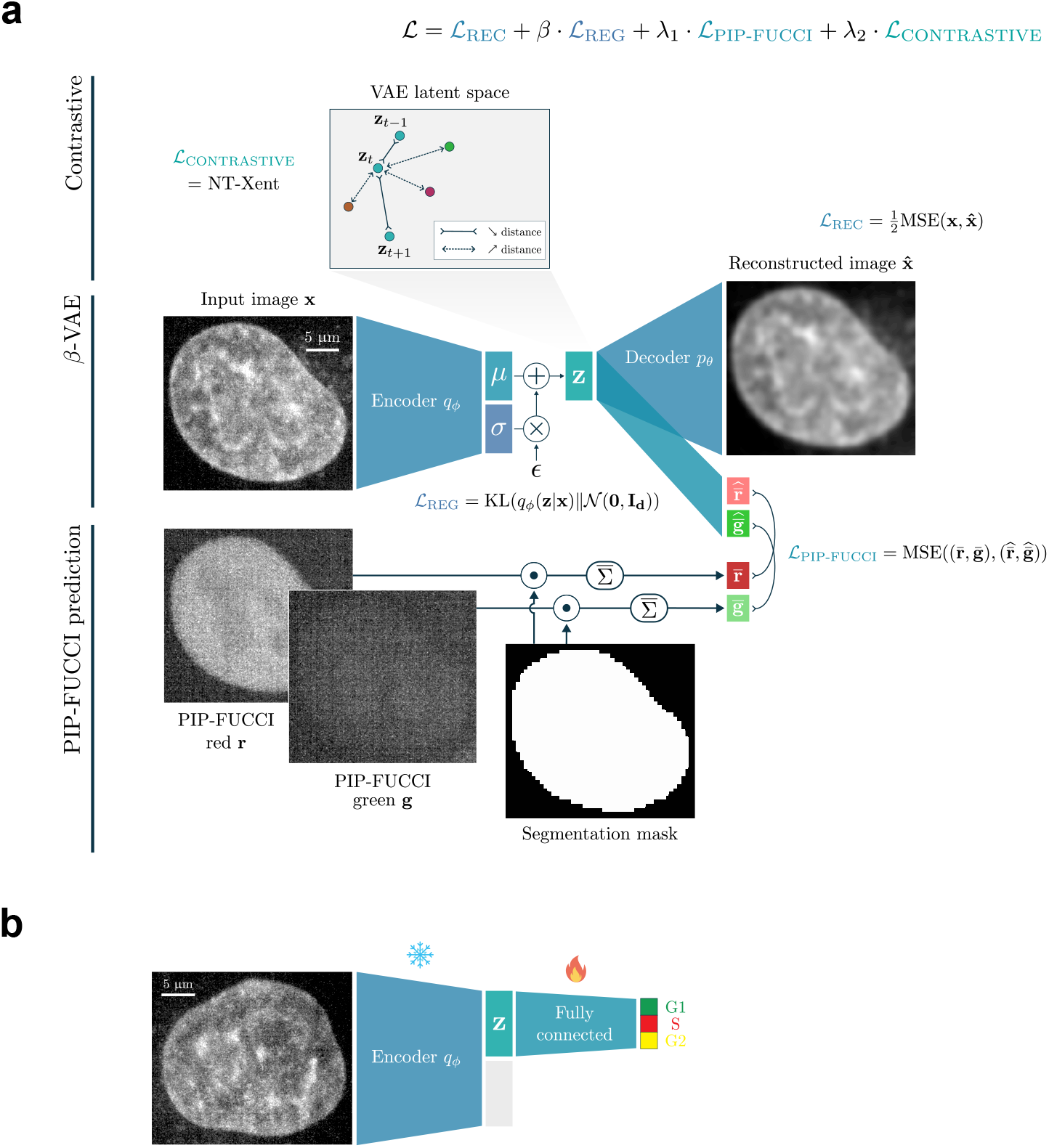
Architecture of Cell-Cycle VAE. (a) Building on the *β*-VAE architecture, we enhance the model by predicting the average intensity of both PIP-FUCCI signals from the latent representation and ensuring temporal consistency through a contrastive loss applied in the latent space. (b) For classification, two fully connected layers are added on top of the frozen CC-VAE encoder to perform supervised training.

Vanilla VAEs [31] compress an original image **x** into its latent representation **z** ∈ ℝ^*d*^ through the encoder *q*_*ϕ*_. The decoder *p*_*θ*_ then uses this latent representation as input to reconstruct the initial image, 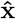. By constraining the latent distribution *q*_*ϕ*_(**z** | **x**) to follow a standard Gaussian 𝒩 (**0, I**_**d**_), VAEs learn a robust and powerful latent representation. Both the encoder and decoder are parameterized by deep neural networks. This allows VAEs to be trained by minimizing the sum of a reconstruction loss, 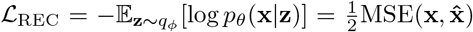, and a regularization loss, ℒ_REG_ = KL(*q*_*ϕ*_(**z** | **x**) ∥ 𝒩 (**0, I**_**d**_)) – where MSE and KL refer to the Mean-Square Error and the Kullback-Leibler divergence, respectively. As [33] introduced a parameter *β* to balance between reconstruction accuracy and latent space regularization, *β*-VAE are trained by minimizing the loss function:

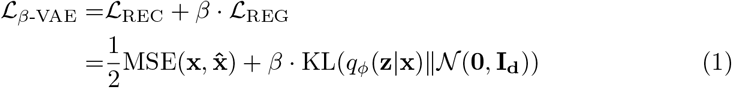

Building on this approach, we aimed to adapt it for the task of learning time-consistent, cell cycle-aware latent representations of cell nuclei.

First, we use the PIP-FUCCI channels during the self-supervised training. Inspired by in-silico labeling strategies described in [34, 35], we utilize our latent representation to predict the average intensity of both PIP-FUCCI signals across the nucleus area. Since the total fluorescent intensities are closely associated with cell cycle phases, this additional task helps structure the latent space to better support downstream cell cycle phase classification.

Second, we exploit the time dependency of our images to enforce time consistency within the latent space. Contrastive learning [36] aims at learning rich image representations by generating multiple views of the same input image using semantic-preserving data augmentations, and train models to group those views together in representation space. If two views come from the same input image (defining a positive pair), their representations should be close; otherwise (negative pair), their representations should be dissimilar and thus more distant. Here, we assume that the nuclei do not drastically change from one time frame to another. We therefore expect that the representations of one nucleus at two consecutive time frames should be relatively close in the latent space. This one-frame shift can therefore be seen as a semantic-preserving image augmentation. Consecutive frames of the same nucleus would be treated as positive pairs, while all other nuclei within the training batch would be treated as negative samples. To encourage such positive pairs to move towards each other and negative pairs to move away from each other, we incorporate the Normalized Temperature-Scaled Cross-Entropy (NT-Xent) loss, as proposed in [37], into our training process. For a positive pair of examples (*i, j*), this loss function writes:

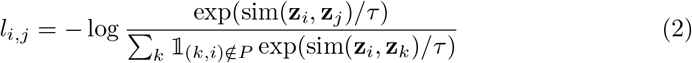

where sim refers to the cosine similarity function, *τ* is the temperature parameter, and *P* is the set of positive pairs within the corresponding training batch. 𝟙_(*k*,*i*)∉ *P*_ is the indicator function evaluating 1 iff (*k, i*) ∉ *P* . The contrastive loss is computed across all positive pairs in the batch: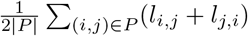.

Our final loss function is thus:

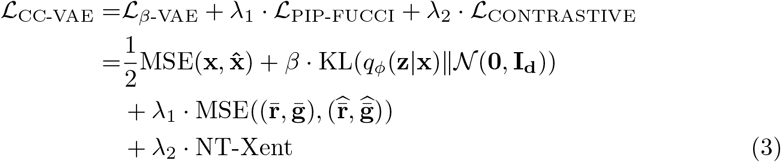

where 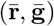 and 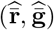 are the true and predicted average PIP-FUCCI mCherry-Gem_1-110_ (red) and PIP-mVenus (green) intensities, respectively.

Following this self-supervised training, we train two fully connected layers on top of the frozen CC-VAE encoder features to predict the cell cycle phase for a given nucleus (see Fig.2b).

### CC-VAE learns rich nucleus representations

CC-VAE allows us to map each nucleus image to a representation in a latent space. Fig.3 shows the UMAP representation of the learned embeddings from a subset of the test set, specifically 276 cell tracks that comprise all three cell cycle phases G1, S and G2/M. Each dot represents a nucleus. In particular, the nuclei presented in Fig.1 are localized in the latent space. Even though the cell cycle information was not used at all during self-supervised training, CC-VAE effectively disentangles the latent space, such that each phase corresponds to coherent regions in the latent space.

**Fig. 3:**
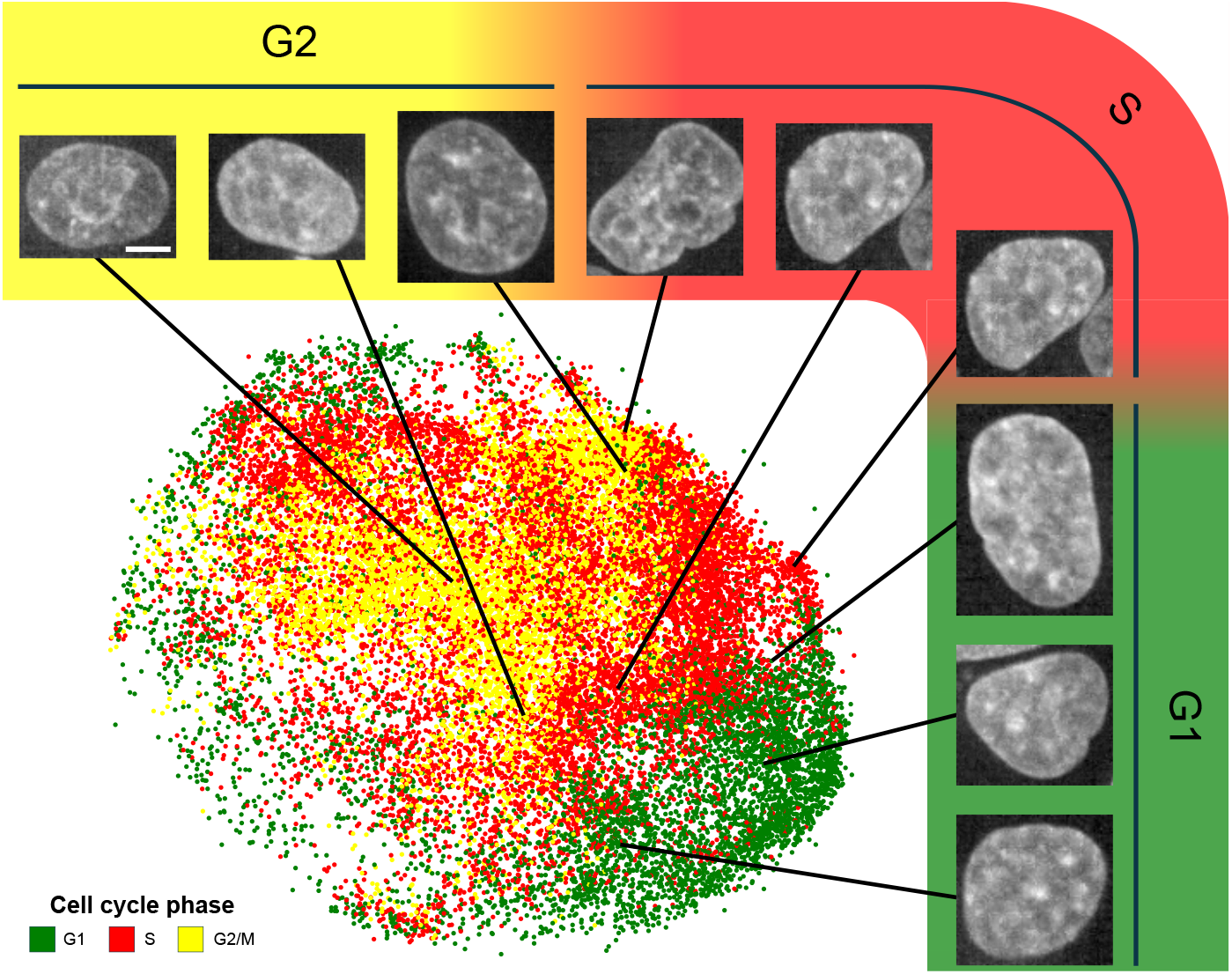
UMAP representation of the nucleus representations learned by Cell-Cycle VAE. Each dot corresponds to a nucleus, colored by cell cycle phase. Scale bar: 5µm.

Fig.4 displays the same UMAP representation, with nuclei colored according to different rules. In Fig.4b, nuclei are colored depending on the time elapsed since mitosis. CC-VAE successfully captures this temporal progression: early-cycle nuclei are positioned at the edges of the latent space, while late-cycle nuclei cluster toward the center. This spatial organization highlights the model’s ability to encode temporal information related to cell cycle progression in its learned representations.

**Fig. 4:**
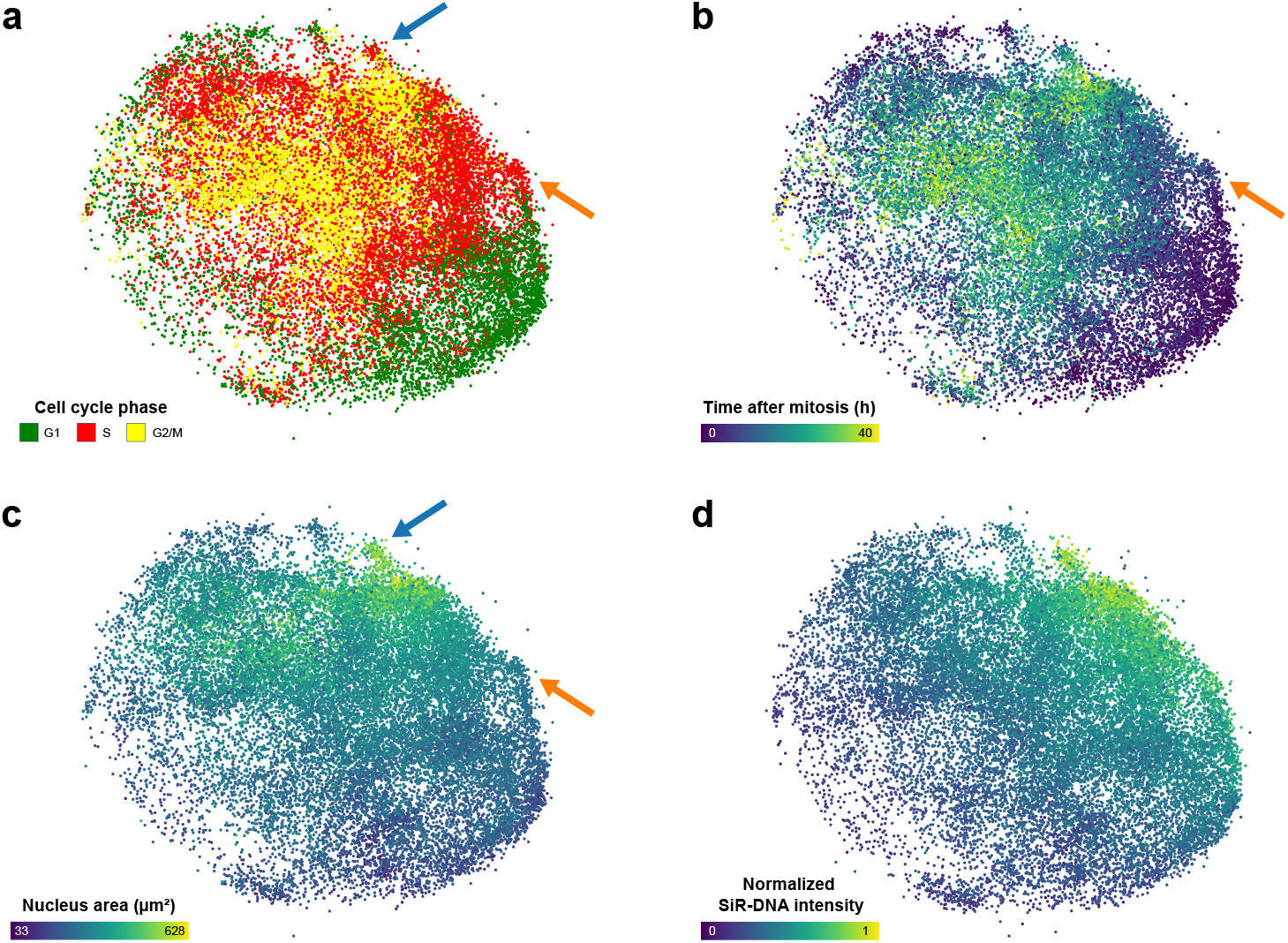
UMAP representation of the nucleus embeddings learned by Cell-Cycle VAE. Each dot corresponds to a nucleus, colored by: (a) cell cycle phase (b) time elapsed since mitosis (c) area (d) normalized SiR-DNA intensity. The blue arrow points to large S-phase nuclei, and the orange arrow points to small early S-phase nuclei. These nuclei could be misclassified as G2/M or G1, respectively, emphasizing the necessity for a classification model that captures subtle phenotypic details beyond basic mor-phological features.

Fig.4c and Fig.4d present the latent space with nuclei colored by their area and total SiR-DNA intensity normalized across all nuclei, respectively. In Fig.4c, larger nuclei locate at the center and top of the latent space, which corresponds to the G2/M phase. This placement aligns with the expected biological progression, as nuclei increase in size throughout the cell cycle, reaching their maximum dimensions during the G2/M phase [29, 30]. In Fig.4d, nuclei with higher SiR-DNA intensity are located on the top-right side of the latent space, corresponding to the S and G2/M phase.

Additionally, our model identifies specific nuclei that would have been misclassified if only area and SiR-DNA intensity were considered. The small cluster of nuclei marked by the blue arrow in Fig.4a and Fig.4c represents large S-phase nuclei, which could be mistaken for G2/M nuclei due to their size. Similarly, nuclei indicated by the orange arrow in Fig.4a, Fig.4b, and Fig.4c correspond to small early S-phase nuclei, which could be incorrectly classified as G1 nuclei. This demonstrates CC-VAE’s ability to capture subtle phenotypic differences beyond basic morphological features.

As our dataset is derived from live-cell imaging, it is possible to compute the latent representation of all nuclei within a specific track, allowing to trace the cell trajectory in the latent space. This representation visualizes the nucleus evolution throughout the cell cycle. Fig.5 displays six distinct nucleus trajectories transitioning from G1 to G2/M. This visualization highlights the temporal consistency achieved by our approach, where nuclei that are phenotypically close (such as the same nucleus in subsequent time frames) are also close in latent space.

**Fig. 5:**
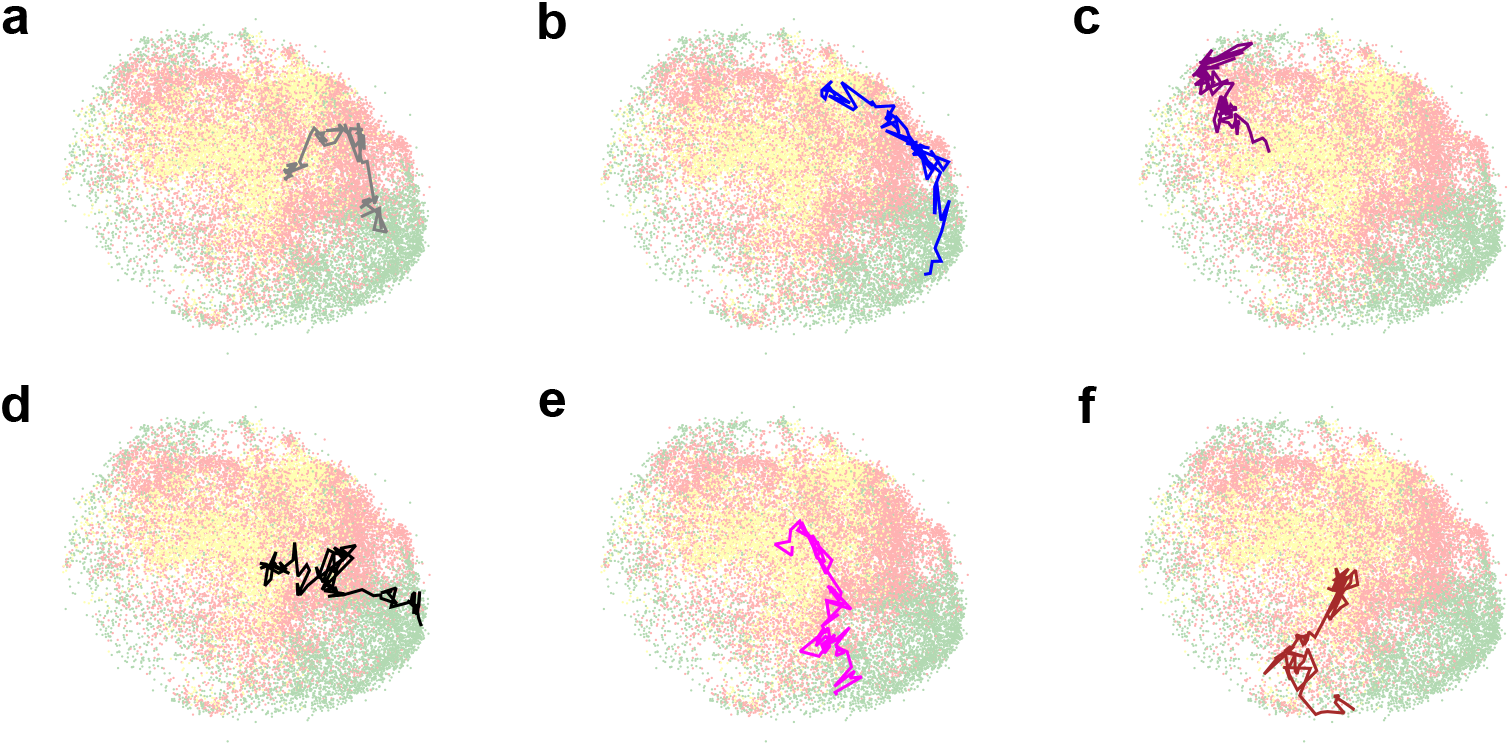
(a-f) Nucleus trajectories in the UMAP representation of the CC-VAE latent space. Each trajectory follows a nucleus over time, from the G1 phase to the G2/M phase.

### CC-VAE classifies cell cycle phases with high accuracy

Given the structure of the latent space, the learned latent representations can be used to train a robust cell cycle phase classifier. Fig.6 shows the averaged confusion matrix from our 10-fold cross-validation dataset (see Methods). We observe good performance with an average recall of 82% across classes (86% for G1, 77% for S and 83% for G2/M) and average precision of 82% (88% for G1, 80% for S and 78% for G2/M).

**Fig. 6:**
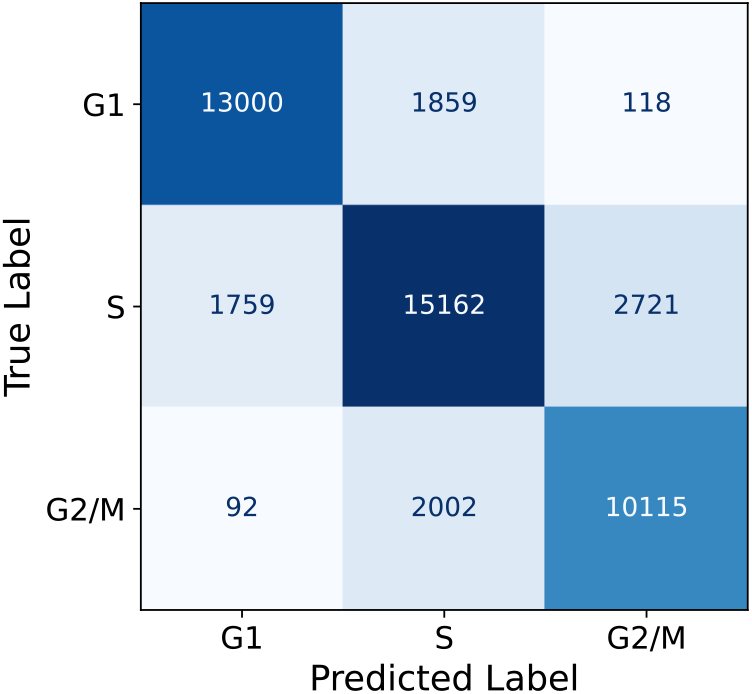
Average confusion matrix from 10-fold cross-validation of CC-VAE.

As expected, misclassifications between G1 and G2/M nuclei are rare (only 2.5% of all errors) due to their distinct features. Although the S phase remains the most challenging to classify due to overlapping boundaries with both G1 (42.3% of errors are G1 vs. S confusions) and G2/M (55.2% of errors were G2/M vs. S confusions), our approach demonstrates reliable performance with 77% recall and 80% precision for S as the most difficult class.

Next, we compared the performance of our model to two alternative strategies for nucleus classification, inspired by related works mentioned above. Since these models could not be used directly due to differences in input modality, we trained them on our dataset to allow for a fair comparison of their relative performances.

First, we trained a Support Vector Machine (SVM) classifier with linear kernel that took as input the nucleus’s area and total SiR-DNA intensity. This baseline corresponds to what was traditionally used as a cell cycle proxy [26]. Next, we trained a CNN with the same backbone as CC-VAE and pretrained on ImageNet, a dataset of natural images [22–25]. Note that all CNN weights were trainable during the classification training, whereas our CC-VAE approach only trains the last two fully connected layers on top of the frozen features.

The methods were evaluated using 2 different metrics.

First, we evaluated time consistency using the top-1 retrieval accuracy, adapted from [38]. This metric assesses the model’s ability to accurately retrieve positive pairs (consecutive frames) within a batch, reflecting the temporal coherence of the learned representations. Specifically, for a positive pair (*i, j*) in a batch, the retrieval is considered successful if 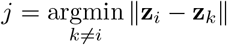, meaning that the closest representation to *i* within the batch (in terms of Euclidean distance) is *j*. The final score is obtained by averaging this result across all possible pairs of all elements in the test set.

Second, we evaluated the cell cycle phase classification performance using macro-averaged accuracy. This metric ensures fair handling of class imbalance by averaging accuracy across all classes. Results are displayed in Table 1.

**Table 1:**
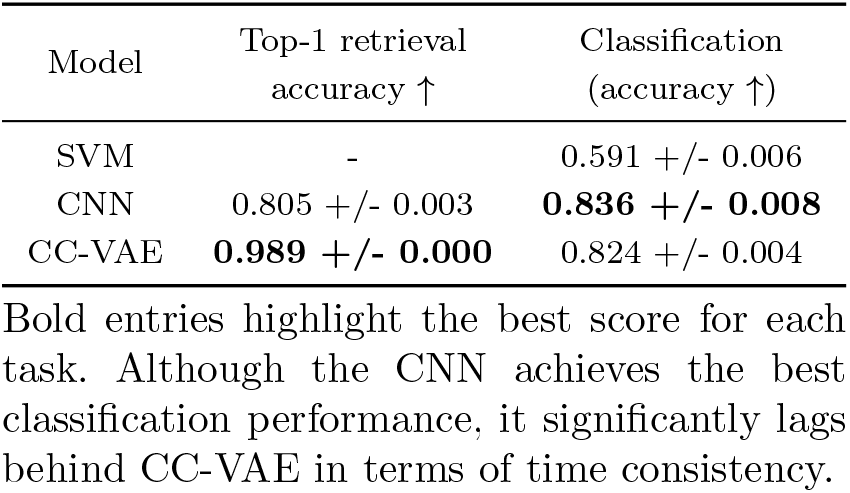
Benchmarking results.

We observe that the CNN trained from scratch slightly outperforms CC-VAE (0.836 +/-0.008 vs 0.824 +/-0.004). However, this approach significantly underperforms in terms of time consistency, as it struggles to achieve the high top-1 retrieval accuracy of CC-VAE (0.805 +/-0.003 vs 0.989 +/-0.000). This quantitative difference in performance is further illustrated by the trajectories of individual nuclei throughout the cell cycle. Fig.7 displays the nucleus trajectories in the CNN latent space, for the same tracks shown in Fig.5. While CC-VAE shows consistent trajectories that allow for smooth tracking of nuclei throughout the cycle, the CNN exhibits highly discontinuous and unstable trajectories, with large oscillations from one extreme of the latent space to the other. Such a lack of temporal consistency makes the CNN representations less reliable and less aligned with biological reality, as they fail to capture the continuous progression of the cell cycle.

**Fig. 7:**
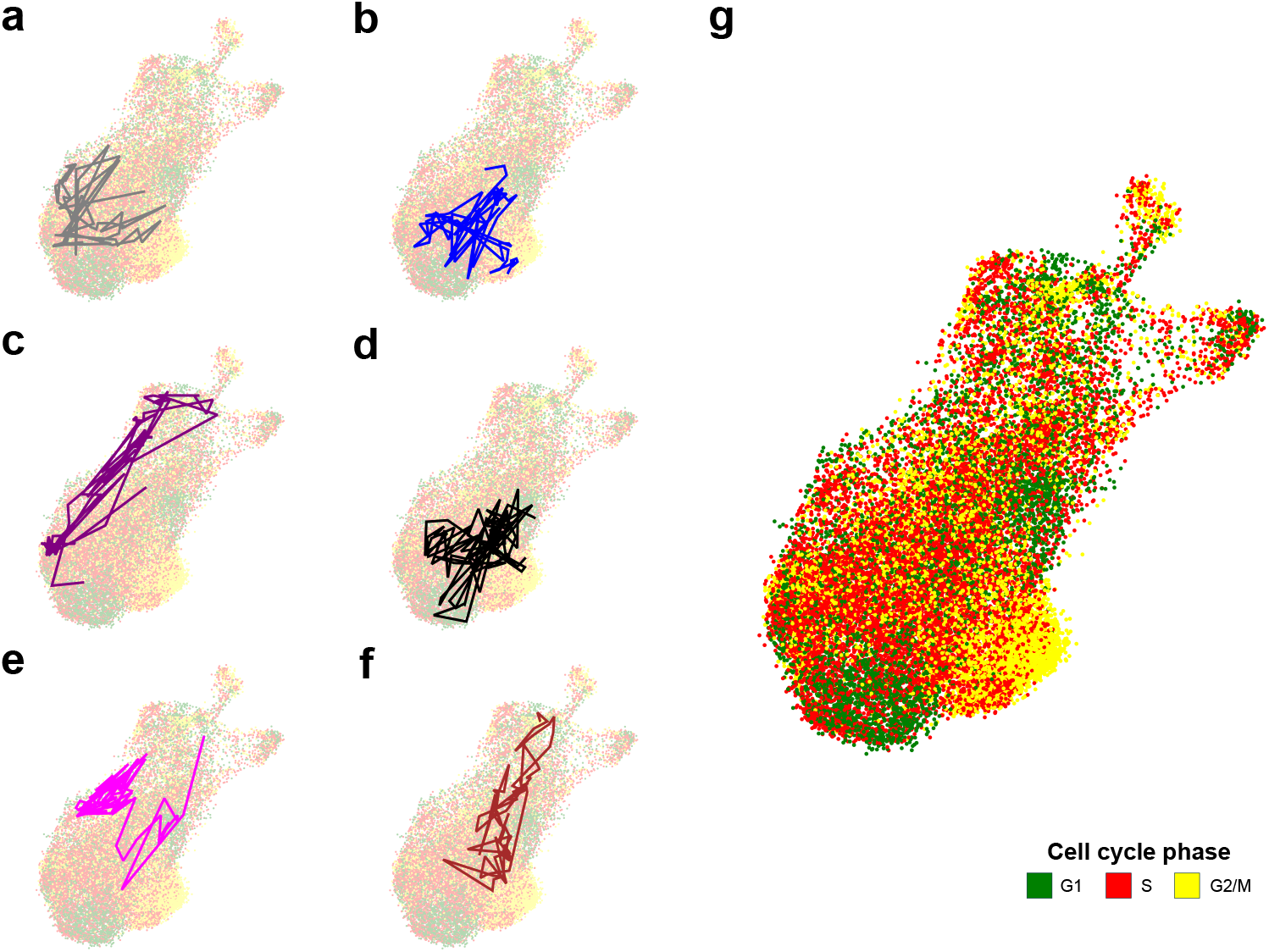
(a-f) Nucleus trajectories in the UMAP representation of the CNN latent space (g). Displayed tracks originate from the same cells as those in the CC-VAE latent space.

### Ablation study

We conducted an ablation study to investigate the impact of the different algorithmic elements. This study compares the performance of different models by successively removing specific elements from the CC-VAE method.

All models share the same backbone architecture. After potential pretraining on a pretext task, the encoder weights are frozen, and only the final two fully connected layers are trained for supervised cell cycle phase classification.

The first model is not pretrained on any task, and its weights are therefore randomly initialized. Next, we evaluated a model pretrained on the standard ImageNet classification task. We then tested a vanilla *β*-VAE, followed by our CC-VAE trained to predict PIP-FUCCI mean intensity as secondary task (ℒ = (ℒ_*β*-VAE_ + *λ*_1_ · ℒ_PIP-FUCCI_). Finally, we assessed our complete model, Cell Cycle-VAE, with (ℒ_CC-VAE_ = ℒ_*β*-VAE_ + *λ*_1_ ·ℒ_PIP-FUCCI_ + *λ*_2_ · ℒ_CONTRASTIVE_.

Additionally to the two metrics presented in the benchmarking, for VAE models, the reconstruction error was assessed using the Structural SIMimlarity (SSIM) metric. SSIM focuses on structural information rather than pixel-wise differences, making it particularly suitable for microscopy image analysis compared to traditional metrics like Mean Squared Error (MSE). Results are displayed in Table 2.

**Table 2:** Ablation study.

| Pretext task | Reconstruction<br>(SSIM ↑) | Top-1 retrieval<br>accuracy ↑ | Classification<br>(accuracy ↑) |
| --- | --- | --- | --- |
| None | - | 0.911 +/- 0.022 | 0.526 +/- 0.008 |
| ImageNet classification | - | 0.790 +/- 0.002 | 0.635 +/- 0.004 |
| β-VAE | **0.655 +/- 0.004** | 0.957 +/- 0.001 | 0.697 +/- 0.008 |
| CC-VAE, $\lambda_2 = 0$ | 0.639 +/- 0.003 | 0.963 +/- 0.001 | 0.819 +/- 0.003 |
| CC-VAE | 0.633 +/- 0.003 | **0.989 +/- 0.000** | **0.824 +/- 0.004** |

Bold entries highlight the best score for each task. While the model pretrained with $\beta$ -VAE achieves the best reconstruction, our proposed method CC-VAE outperforms other methods both on time regularization and cell cycle classification.

The *β*-VAE achieves the best performance in reconstruction, which is expected since the model does not have to perform the additional tasks of Cell Cycle-VAE, leading to fewer constraints on the model. While the two Cell Cycle-VAEs perform quite similarly in terms of classification accuracy, the addition of the time consistency regularization term in the loss enhances the time regularization of the latent space. This regularization helps learning features that are more aligned with the phenotypic characteristics of the nuclei.

## Discussion

We present Cell Cycle Variational Auto-Encoder (CC-VAE), a novel deep self-supervised method for determining the cycle phase of individual cells based on a DNA fluorescence marker. CC-VAE can serve as a powerful tool for cell stratification according to their cycle phase, enabling for example the study of new connections between the cell cycle and patterns of RNA or protein localization.

For this, we propose variational auto-encoders, that provide generic and powerful latent space representations. We propose two auxiliary tasks, prediction of the cell-cycle dependent PIP-FUCCI fluorescent intensities and a novel time-consistency constraint. Together, they structure the latent space to group nuclei according to their cell cycle phase while preserving phenotypic continuity ensuring that nearby nuclei in the latent space are also phenotypically close. A lightweight classifier trained on the frozen latent representations achieves an average classification accuracy of 82.4%. All code and the trained networks are made freely available at GitHub, https://github.com/15bonte/cell_cycle_classification and HuggingFace, https://huggingface.co/thomas-bonte/cell_cycle_classification, respectively.

Notably, our strategy optimizes relatively few parameters specifically for classification. While training a full CNN end-to-end for classification can yield slightly higher accuracy, we show that this comes at the cost of temporal and phenotypic consistency. Temporal consistency is particularly important, as it reflects the inherently continuous nature of the cell cycle, even though we discretize it into three main phases in this work. Relying on latent representations that do not preserve biological distances can be misleading, as they might suggest phenotypic discontinuities that do not correspond to a biological reality. Variational auto-encoders provide a natural mechanism to add biologically meaningful pretext tasks and thus impose a biological structure to the latent space.

Another important aspect of the study is the overall annotation strategy. We found that tracking PIP-FUCCI over time is essential to reliably identify cell cycle phases due to high variability in marker intensity between individual cells. While the intensity time series allow for a reliable cell cycle assessment, static snapshots may lead to label noise that propagates through the learning process. Here, we have carefully annotated a large number of cells over time, leading to a dataset of over 600,000 annotated nuclei. This dataset is made freely available on BioImage Archive at https://www.ebi.ac.uk/biostudies/bioimages/studies/S-BIAD1659 to support further community development and benchmarking.

Our current approach is not free of limitations. First, we can expect some improvements by incorporating additional label-free modalities, such as bright-field or phase-contrast imaging, providing complementary phenotypic information missing from DNA fluorescence markers, and thus potentially improving both cell cycle phase prediction and overall phenotypic consistency of the latent space. This enhancement would be particularly valuable for biologists as these modalities are common, label-free, and already frequently used in daily experiments. Furthermore, the current approach is still vulnerable to domain shifts induced by the use of other DNA markers, thus limiting direct out-of-the-box use of the trained networks. However, since SiR-DNA and Hoechst are derived from the same molecule [32], and both SiR-DNA and DAPI intercalate into AT-rich regions of DNA [32, 39], it may be possible to use domain adaptation techniques to adapt our networks to these markers. This adaptation could help democratize CC-VAE, making it a more accessible and practical tool for biologists. Finally, CC-VAE enhances the *β*-VAE architecture by integrating in-silico labeling and time-contrastive pretext tasks, in order to structure the latent space for cell cycle disentanglement. However, alternative pretext tasks could have been explored. Notably, Fig.4b and Fig.8 suggest – as expected – a strong correlation between cell cycle phases and the time elapsed after mitosis. Predicting this temporal information could serve as a powerful pretext task for cell cycle phase classification. Yet, this approach is not feasible in our case, as most tracks do not span the entire cell cycle but only partial segments. As a result, the exact onset of the cycle remains unknown for most nuclei, preventing access to precise elapsed time since mitosis. However, this method could be worth exploring on a data set composed exclusively of tracks that span the entire cell cycle.

**Fig. 8:**
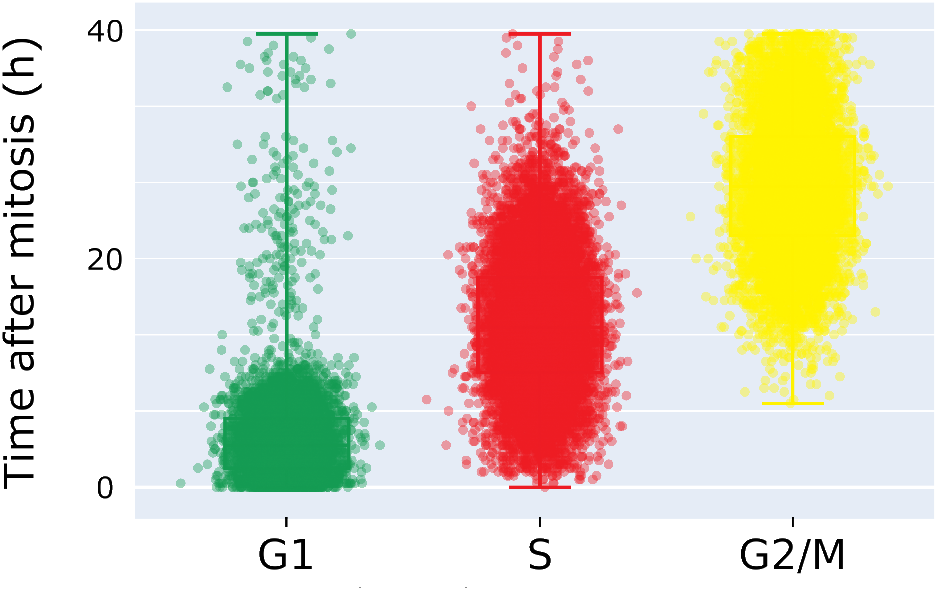
Time elapsed after mitosis (hours). Each dot corresponds to a nucleus from the same subset of the test set as in Fig.3.

## Conclusions

Here, we introduced CC-VAE, a powerful self-supervised-based method for accurately predicting the cell cycle phase from DNA marker microscopy images. It improves the *β*-VAE architecture by incorporating in-silico labeling and time-contrastive additional tasks, enabling the model to capture subtle phenotypic information related to cell cycle progression. Leveraging this rich latent encoding, CC-VAE achieves high accuracy and robustness in cell cycle phase classification. Additionally, we provide a novel dataset containing over 600,000 labeled nucleus images, which we believe will be a valuable resource for ongoing and future research in the field.

## Methods

### Variational Auto-Encoders

Variational Auto-Encoders (VAE, [31]) belong to the family of self-supervised generative models. Compared to vanilla auto-encoders, they provide better latent representations by introducing regularization, both at the local and global level. In the following, bold symbols (e.g., **x**) denote vector quantities.

Let **x** ∈ ℝ^*D*^ be a set of observable variables deriving from an unknown distribution. VAE assume that there exist latent variables **z** ∈ ℝ^*d*^, **z** being a latent representation of **x** with *D* ≫ *d*, i.e. the latent space is of much smaller dimension than the original space. The generative model writes **z** ∼ *p*_**z**_(**z**) ; **x** ∼ *p*_*θ*_(**x**|**z**), with *p*_*θ*_(**x**|**z**) taken as a parametric distribution. Therefore, the marginal likelihood *p*_*θ*_(**x**) is given by:

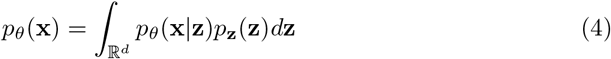

Since the integral is taken over the entire latent space, it is most of the time computationally intractable – meaning that the computation complexity of any approach for evaluating this integral is exponential. So does the posterior distribution *p*_*θ*_(**z**|**x**), which is then approximated by a parametric distribution *q*_*ϕ*_(**z**|**x**) through variational inference [40]. *q*_*ϕ*_(**z** | **x**) and *p*_*θ*_(**x** | **z**) are refered to as the *encoder* and the *decoder*, respectively, whose parameters are given by deep neural networks. Importance sampling enables to define an unbiased estimate 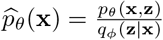 of the marginal likelihood *p*_*θ*_(**x**), and to obtain a lower bound for the marginal log-likelihood through Jensen’s inequality:

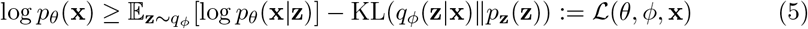

ℒ (*θ, ϕ*, **x**) is called the Evidence Lower BOund (ELBO). Since we aim at maximizing the marginal log-likelihood log *p*_*θ*_(**x**) (as the probability of the model itself), the ELBO can be considered as a good proxy to maximize during training. In vanilla VAEs, the prior *p*_**z**_(**z**) is chosen as a standard Gaussian distribution 𝒩 (**0, I**_**d**_).

### Live cell imaging

Hela Kyoto cells stably expressing Cdt1(1-17aa)-HA-mVenus and mCherry-Geminin(1-110aa) were cultured in 10 cm dishes in DMEM + 10%FBS + 1% PenStep at 37°C with 5% CO_2_. For imaging, cells were cultured on a 96-well phenoplate (Revvity). Culture media was aspirated and cells were incubated with 100 nm SiR-DNA Cy5 (Spirochrome) for 4h at 37°C. The media was then replaced with imaging media pre-warmed at 37°C (DMEM fluorobrite, 1% P/S, 10% FBS, glutamax, 100 nm SiR DNA) at least 2h before imaging.

One day after seeding, time-lapse microscopy was performed via live-cell fluorescence imaging using an Opera Phenix™ Plus High-Content Screening System (Perkin Elmer) with a 63X water immersion objective (1.15 NA). Imaging lasted for 40 hours, capturing images every 20 minutes. A total of 159 fields-of-view were acquired, with 5 focal planes per field spaced 1 µm apart. 3 channels (488 nm, 561 nm and 640 nm) were acquired, each with 20% laser power and 100 ms exposure time. The temperature and CO_2_ level were maintained at 37°C and 5%, respectively. One pixel corresponds to 0.102 µm.

### Image preprocessing

Nucleus segmentation was performed on each frame independently, using Cellpose [41] applied to the SiR-DNA channel. We used the Cellpose interface to conduct human-in-the-loop fine-tuning of the default Cellpose segmentation model. This involved using 20 frames randomly selected from our dataset. We retained Cellpose’s default training parameters: a learning rate of 0.1, weight decay of 0.0001, and 100 epochs. Segmentation was performed with the flow threshold set to 0.4 and nucleus probability threshold set to 0. The nucleus diameter is automatically calculated by Cellpose and set to 173.90 px, i.e. 17.74 µm.

The tracking is performed using the particle tracking algorithm developed by [42] and implemented in TrackMate [43]. The maximum distance for tracking is set to 200 px, i.e. 20.4 µm. Track merging, track splitting and gap closing are not permitted. Across the dataset, a total of 15,026 nucleus tracks were identified, encompassing 982,332 single nucleus images.

The nuclei were labeled on the basis of the temporal evolution of their PIP-FUCCI intensities across their area. 217,465 nuclei were labeled as G1, 257,984 were labeled as S, and 160,855 were labeled as G2/M. Additionally, 346,028 nuclei remained unlabeled.

### Deep learning training

The 159 fields-of-view in our dataset are divided into 10-fold cross-validation sets, ensuring that nuclei from the same field-of-view are grouped within the same set. Each fold includes 80% of the data for training, 10% for validation, and the same 10% for testing.

Our encoder is a standard ResNet18 with 11.4M parameters. The decoder, designed to mirror the encoder’s structure, is symmetrical. The self-supervised model was trained for 10 epochs with a learning rate of 1e-4 and a batch size of 128. One epoch takes on average 2 hours on a P100 GPU. We used data augmentations including rotation, horizontal flip, and vertical flip. Early stopping was employed to select the best model based on validation performance. The implementation of the VAE-based model relies on Pythae [44].

The average intensities of PIP-FUCCI are predicted using a single linear layer applied to the latent representation **z**. The contrastive NT-Xent loss is applied in a manner similar to the approach described in [37], with the key difference being that the only augmentation introduced in our method is a one-frame shift. We retained the default temperature parameter value of 0.7.

For supervised classification training, we train two fully connected layers (67k parameters) on top of the frozen encoder features to classify the nucleus images into one of three phases: G1, S, or G2/M. Training was performed during 1 epoch using a learning rate of 1e-4 and a batch size set to 64, with the cross-entropy loss function used to optimize classification accuracy. Early stopping was employed to select the best model based on validation performance.

To improve classification accuracy, we performed a grid search to select optimal values for *β*, the self-supervised learning rate, the latent dimension, *λ*_1_ and *λ*_2_. To limit training time, this grid search was performed on 5% of the train and validation data. Additionally, to reduce the number of combinations, we first considered the three parameters used in the standard *β*-VAE (*β*, learning rate, latent dimension), followed by *λ*_1_, and finally *λ*_2_. *β* was chosen in {0.01, 0.1, 1} as [44] suggests that selecting *β* > 1 can negatively impact classification accuracy. The learning rate was chosen in {5e − 5, 1e − 4, 5e − 4} and the latent dimension was chosen in {64, 128, 256, 512} . The optimal parameters for *β*-VAE were found to be *β* = 0.01, a latent dimension of 256 and a learning rate 1e-4. *λ*_1_ and *λ*_2_ are chosen in {10, 100, 1e3, 1e4, 1e5} and {1, 10, 100, 1e3}, respectively. These sets were chosen based on the initial ratio between the *β*-VAE loss and the additional losses. The optimal parameters were found to be *λ*_1_ = 1e4 and *λ*_2_ = 1e2.

The CNN presented in Table 1 is a ResNet18 pretrained on ImageNet classification and trained using the same parameters as above, except that the number of epochs was increased from 1 to 10 to ensure effective classification learning.

## Declarations

### Ethics approval and consent to participate

Not applicable.

### Consent for publication

Not applicable.

### Availability of data and materials

The datasets generated and analysed during the current study are available in the BioImage Archive repository, https://www.ebi.ac.uk/biostudies/bioimages/studies/S-BIAD1659.

The Python code is available via GitHub, https://github.com/15bonte/cell_cycle_classification.

The tuned models are available via HuggingFace, https://huggingface.co/thomas-bonte/cell_cycle_classification.

### Authors’ contributions

T.W. and E.B. supervised the project. O.P. performed all imaging experiments, under the supervision of E.B. T.B. annotated the dataset. T.B. designed and implemented Cell Cycle-VAE in Python, and analyzed the results with input from all the authors. T.B. and T.W. wrote the manuscript, T.B. prepared all figures. A.S., D.W., F.M. and F.S. contributed ideas and suggested experimental directions. All authors reviewed and approved the final manuscript.

## Acknowledgements

Not applicable.

